# A unified characterization of population structure and relatedness

**DOI:** 10.1101/088385

**Authors:** Bruce S. Weir, Jérôme Goudet

## Abstract

Many population genetic activities, ranging from evolutionary studies to association mapping to forensic identification, rely on appropriate estimates of population structure or relatedness. All applications require recognition that quantities with an underlying meaning of allelic identity by descent are not defined in an absolute sense, but instead are made “relative to” some set of alleles other than the target set. The early Weir and Cockerham *F_ST_* estimate made explicit that the reference set of alleles was across independent populations. Standard kinship estimates have an implicit assumption that pairs of individuals in a study sample, other than the target pair, are unrelated, whereas other estimates assume alleles within individuals are not identical by descent. However, populations lose independence when there is migration between them, and when individuals in a study are related it is difficult to see how they can also be non-inbred. We have therefore re-cast our treatments of population structure, relatedness and inbreeding to make explicit that the parameters of interest involve differences of probabilities of identity by descent in the target and the reference sets of alleles and so can be negative. We take the reference set to be for the population from which study individuals have been sampled. We provide simple moment estimates of these parameters, phrased in terms of allele matching within and between individuals for relatedness and inbreeding, or within and between populations for population structure. A multi-level hierarchy of alleles within individuals, alleles between individuals within populations, and alleles between populations allows a unified treatment of relatedness and population structure. Our new estimates appear to be sensitive to rare or private variants, to give indications of the effects of natural selection, and to be appropriate for use in association studies.

## Introduction

We offer here a unified treatment of relatedness and population structure with an underlying framework of alleles being identical by descent, ibd. We follow Thompson (2013) in regarding ibd for a set of alleles as being relative to some other, reference, set: “There is no absolute measure of ibd: ibd is always relative to some reference population.” In other words, ibd implies a reference point and ibd status there is often implicitly assumed to be zero.

A function of ibd of particular interest to us is *F*_*ST*_, which we will show below depends on ibd of pairs of alleles within populations relative to that for pairs of alleles from different populations. The uses of estimates of this quantity are widespread, and here we note a recent discussion by McTavish and Hillis (2015) of the effects of SNP ascertainment, SNP array vs whole-genome sequencing, on inferences about population history. These authors used “pairwise *F*_*ST*_ for all pairs of populations using Weir and Cockerham’s method.” We suggest that a more informative analysis may result from our population-specific *F*_*ST*_ estimates (Weir and Hill, 2002; Weir et al., 2005; Browning and Weir, 2010). Several authors (e.g. Balding and Nichols, 1995; Shriver et al., 2004; Beaumont and Balding, 2004; Gaggiotti and Foll, 2010) have discussed the advantages of working with population-specific *F*_*ST*_ values instead of single values for a set of populations or of values for each pair of populations. We show below that the usual global *F*_*ST*_ measure can be regarded as an unweighted average of population-specific values, and because it is an average it collapses the variation detectable among populations that can indicate the effects of past selection (Weir et al., 2005). The usual measure can otherwise diminish signals of population history and this diminution has become more pronounced as genetic marker data have become richer and real differences among populations have become more evident. As Astle and Balding (2009) noted “population structure and [cryptic] relatedness are different aspects of a single confounder: the unobserved pedigree defining the (often distant) relationships among the study subjects.” A similar point was made by Kang et al. (2010): “The presence of related individuals within a study sample results in sample structure, a term that encompasses population stratification and hidden relatedness.” Our goal is to provide a unified approach to charactering population structure and individual relatedness and inbreeding, both in terms of the underlying parameters and of the methods of estimation.

A consideration of “genetic sampling” (Weir, 1996) makes it clear that population mean ibd for alleles in a single population, or ibd for alleles in a single individual, at one point in time cannot be estimated only from data for that population or that individual as there would be no information about variation over replications of the descent paths from past to present. We might regard multiple loci as providing replication of the genetic sampling process or we might collect data from multiple populations. An exception is when allele frequencies and ibd status in the reference population are assumed known, as is implied for standard methods for estimating relatedness and inbreeding (e.g. Ritland, 1996; Wang, 2014; Purcell et al., 2006; Yang et al., 2011). If, instead, these methods make use of frequencies from a sample of individuals they are providing estimates of the inbreeding or coancestry ibd measures relative to those measures for individuals in the whole sample. This point was also made by Yu et al. (2006) who spoke of “adjusting the probability of identity by state between two individuals with the average probability of identity by state between random individuals” in order to address identity by descent. Other relatedness estimation methods that do not use allele frequencies (e.g. KING-robust, Manichaikul et al., 2010) are estimating ibd between individuals (coancestry) relative to that within individuals (inbreeding: assumed zero for KING-robust).

For both population structure and relatedness we propose the use of allelic matching proportions within and between individuals or populations in order to characterize ibd for an individual or a population relative to a reference set of ibd values. We use allele matching rather than heterozygosity (Nei, 1973) or components of variance (Weir and Cockerham, 1984: hereafter WC84) although the distinction is more semantic than real. Our present treatment also differs from that in WC84 by using unweighted averages of statistics over populations instead of the weighted averages that were more appropriate for the WC84 model of independent populations.

The size of current genetic studies requires computationally feasible methods for estimating relatedness between all pairs of individuals, potentially 5 billion pairs for the TOPMed project (http://www.nhlbiwgs.org). The scale of the task may well rule out maximum likelihood approaches (e.g. Thompson, 1975; Ritland, 1996; Milligan, 2003) and Bayesian methods (e.g. Gaggiotti and Foll, 2010). Moment estimates seem still to be relevant and will be presented here.

## Materials and Methods

### Parameter Values

We write the probability of a pair of alleles being ibd as *θ* without specifying the reference set of alleles. Subscripts will identify the populations or individuals from which the alleles are drawn. Subscript *W* will indicate an average over sets of alleles within individuals or populations, and subscript *B* the average over pairs of distinct individuals or populations.

#### Pairs of Individuals

The coancestry coefficient *θ*_*XY*_ for individuals *X*, *Y* is the probability an allele taken randomly from *X* is ibd to one taken randomly from *Y*. If individual *A* is ancestral to both *X* and *Y*, and if there are n individuals in the pedigree path joining *X* to *Y* through *A*, then 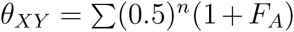 where *F*_*A*_ is the inbreeding coefficient of A and the sum is over all ancestors *A* and all paths joining *X* to *Y* through *A*. The coancestry of *X* with itself is *θ*_*XX*_ = (1 + *F*_*X*_)/2.

#### Pairs of Populations

For populations *i*, *i*′ the quantity *θ*_*ii*′_ is the probability of ibd for an allele from *i* and one from *i*′. The two populations may be the same, *i* = *i*′. The simplest evolutionary scenario is for finite populations of constant size, subject only to genetic drift. For two populations of sizes *N*_1_, *N*_2_ with a common ancestral population *t* discrete generations in the past, the current within-population values *θ*_11_, *θ*_22_ and the between-population value *θ*_12_ are

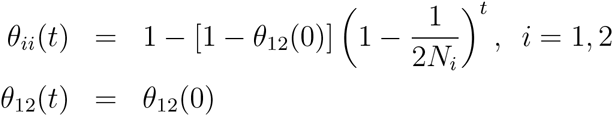

The between-population ibd probability *θ*_12_(*t*) at present is the same as it was, *θ*_12_(0), in the common ancestral population. As we do throughout this discussion, we introduce quantities *β* that measure ibd for a target pair of alleles relative to that in a convenient comparison set, here alleles between the pair of current populations (also, in this case, the common ancestral population):

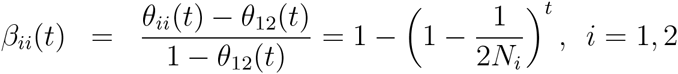

By considering ibd relative to that between populations, we avoid having to know the value in the ancestral population or even having to specify that ancestral population. The two population-specific values *β*_*ii*_(*t*) differ if the two populations have different sizes. We often wish to work with the averages *θ*_*W*_ (*t*) = [*θ*_11_(*t*) + *θ*_22_(*t*)]/2, *θ*_*B*_(*t*) = *θ*_12_(*t*) and we write *β*_*W*_ (*t*) = [*β*_11_(*t*) + *β*_22_(*t*)]/2:

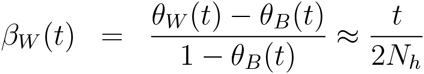

providing *N*_1_, *N*_2_, *t* are all large, and *N*_*h*_ is the harmonic mean of the population sizes. From now on we will regard *β*_*W*_ as the parametric form of *F*_*ST*_, and Reynolds et al. (1983) showed that *F*_*ST*_ serves as a measure of population distance under the pure drift model. The formulation *F*_*ST*_ = (*θ*_*W*_ − *θ*_*B*_)/(1 − *θ*_*B*_) makes explicit that *F*_*ST*_ is a measure of ibd within populations relative to ibd between *pairs* of populations.

#### Drift, Mutation and Migration

Non-trivial equilibria for two populations drifting apart are obtained when there is mutation, and we illustrate some aspects of our population-specific approach by considering the case of two populations exchanging alleles each generation when there is infinite-alleles mutation. The transition equations for *θ*_1_, *θ*_2_, *θ*_12_, extending those given by Maruyama (1970), are:

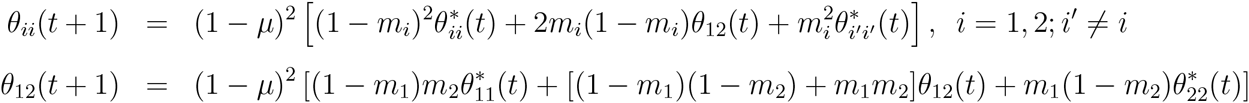

where 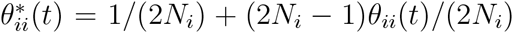, the mutation rate is *µ* and population *i* : *i* = 1, 2 receives a fraction *m*_*i*_ of its alleles each generation from population *i*′ : *i*′ ≠ *i*. A consequence of these equations is that *θ*_11_(*t*) + *θ*_22_(*t*) ≥ 2*θ*_12_(*t*), or that *θ*_*W*_ ≥ *θ*_*B*_ and so *β*_*W*_ = *F*_*ST*_ is positive. However, it is not necessary that each of *θ*_11_, *θ*_22_ exceeds *θ*_12_ and in Figure 1, second row, we show that mutation leads to equilibrium values of *θ*_*ii*_ different from 1, and in Figure 1, third row that migration can lead to cases where *θ*_11_ > *θ*_12_ > *θ*_22_. In the absence of migration, mutation drives *θ*_12_ and *β*_12_ to zero, so that *θ*_*ii*_ = *β*_*ii*_ are both positive. In Figure 2 we show the region in the space of *N*_1_, *m*_1_ values where *β*_11_ ≤ 0 ≤ *β*_22_ for fixed *N*_2_, *m*_2_, *µ*. Averaging over the two *β*_*ii*_’s to work with *F*_*ST*_ hides this potential difference in sign of the *β*_*ii*_’s.

**Figure 1.**
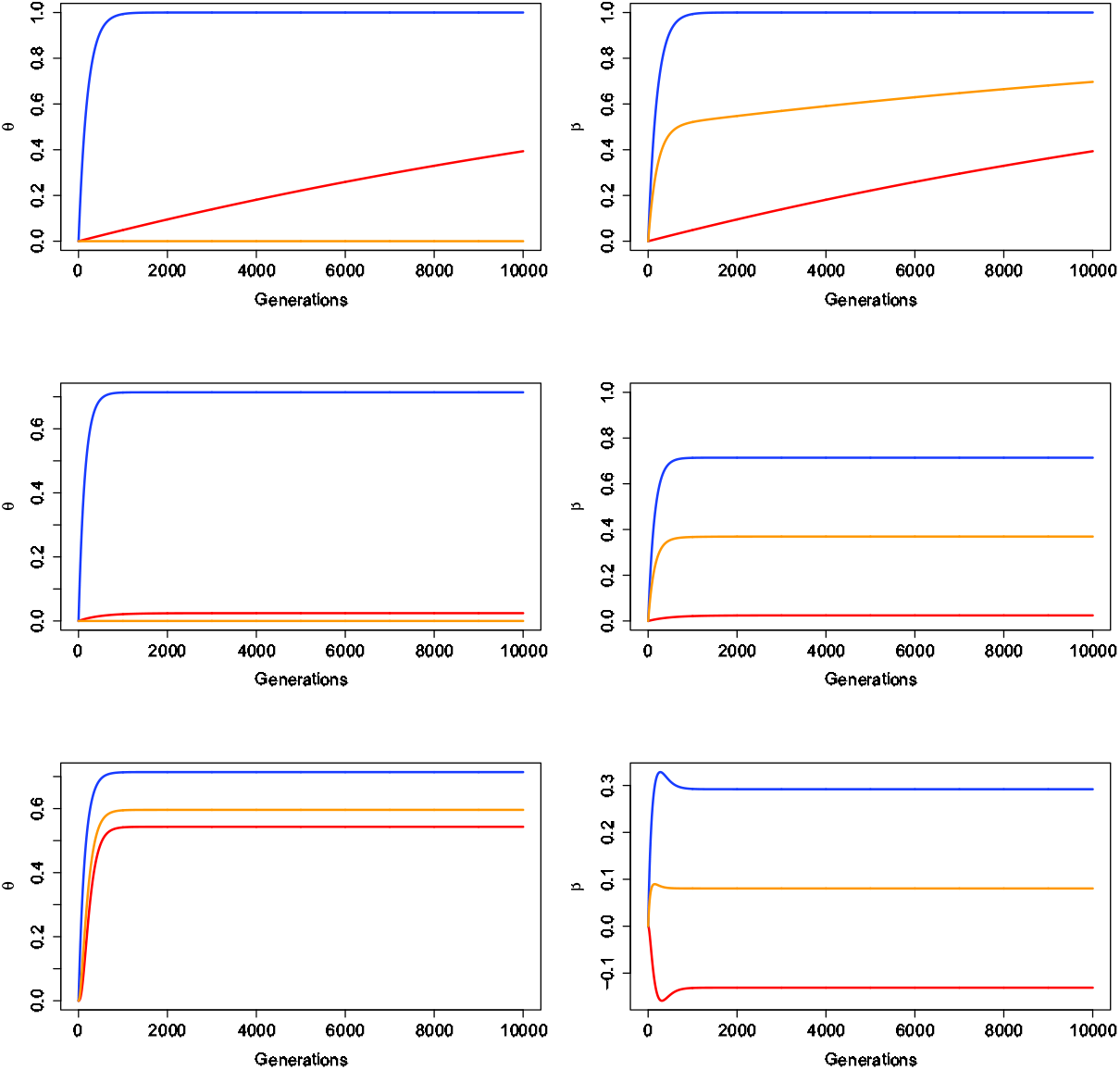
Effects of Drift, Mutation and Migration. For all panels, *N*_1_ = 10000, *N*_2_ = 100 First row: drift only (no mutation nor migration). *θ*_1_, *θ*_2_’s and *β*’s tend to 1,*θ*_12_ = 0.000. Second row: Drift and Mutation *µ* = 10^−3^, *m*_1_ = *m*_2_ = 0. *θ*’s and *β*’s have positive limits less than 1. At equilibrium, *θ*_1_ = 0.024, *θ*_2_ = 0.714, *θ*_12_ = 0.000, *β*_1_ = 0.024, *β*_2_ = 0.714, *β*_*W*_ = 0.369. Third Row: Drift. Mutation and Migration. *µ* = 10^−3^, *m*_1_ = 10^−2^, *m*_2_ = 0. *θ*’s positive and less than 1, *β*_*W*_ is positive but *β*_*ii*_’s may be negative. At equilibrium, *θ*_1_ = 0.543, *θ*_2_ = 0.714, *θ*_12_ = 0.596, *β*_1_ = −0.131, *β*_2_ = 0.292, *β*_*W*_ = 0.080.

**Figure 2.**
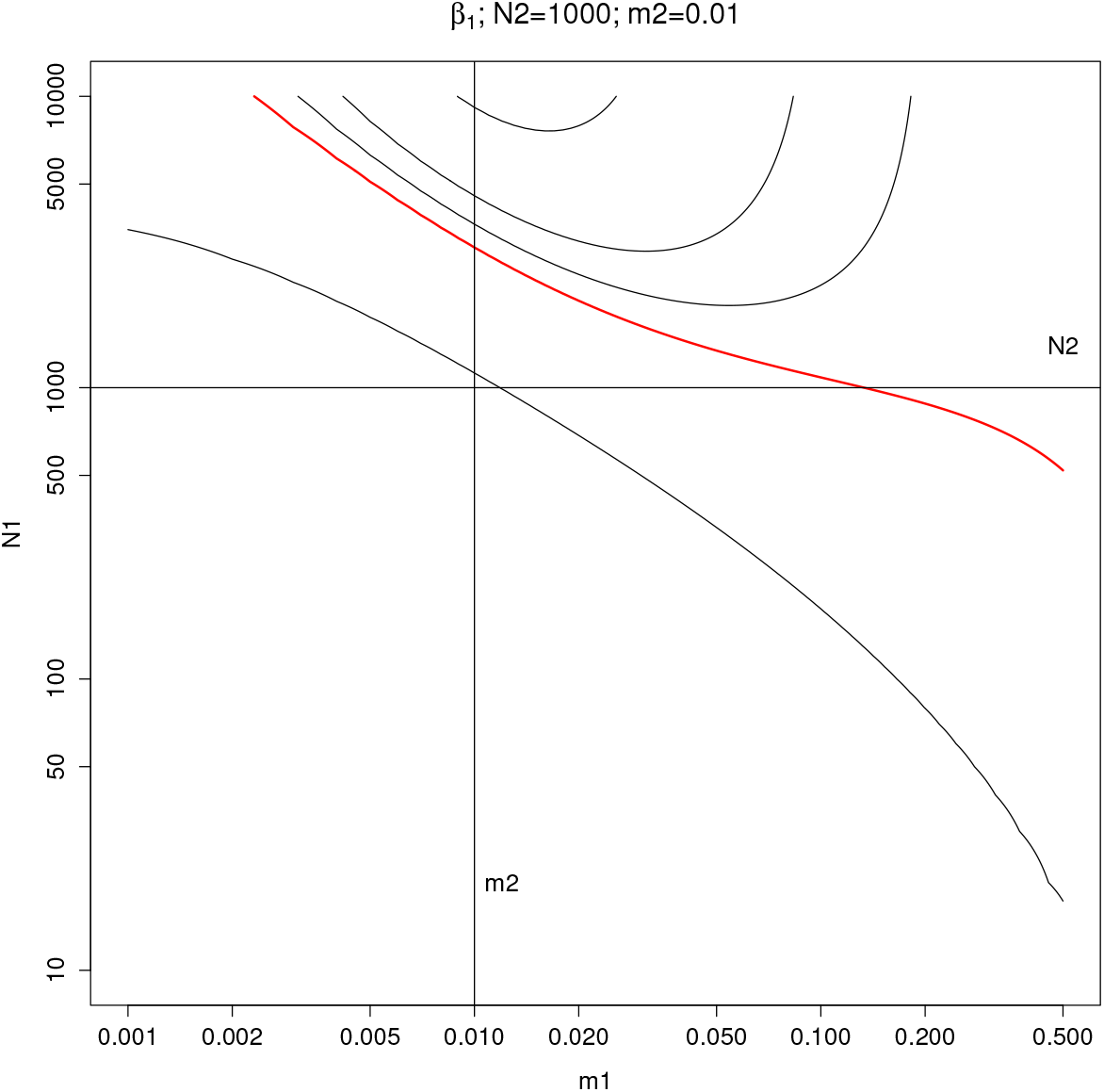
The region above and to the right of the red line has equilibrium values of *θ*_1_ ≤ *θ*_12_ ≤ *θ*_2_, i.e. *β*_1_ ≤ 0 ≤ *β*_2_. In that region a pair of alleles within population 1 has a smaller probability of ibd than does an allele from population 1 with an allele from population 2.

#### Actual vs Predicted

*θ* The probabilities of ibd calculated from path-counting methods for pedigrees of individuals or from transition equations for populations can be regarded as the expected values, over evolutionary replicates, of the actual identity status of a pair of alleles. We have previously discussed the variation of actual identity about the predicted value (Hill and Weir, 2011, 2012). The variance of the actual ibd measure 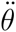 for two alleles is Δ − *θ*^2^ (Cockerham and Weir, 1983), where Δ is the joint probability of ibd for each of two pairs of alleles. The coefficient of variation of the actual coancestry 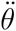 for two individuals is greater than 1 for individuals with predicted coancestry *θ* less than 0.125 and it increases as the degree of relationship decreases. The implication of this is that, for a particular pair of populations or individuals, estimated values may not match those expected from pedigrees or transition equations. Evaluation of estimation procedures should, therefore, be performed over many replicate pairs.

#### Inbred Populations

The discussion so far has implicitly assumed Hardy-Weinberg equilibrium within populations and no need to differentiate pairs of alleles within individuals from those between individuals in the same population. We can relax that assumption. Two alleles taken at random from population *i* may be from the same or different individuals. If the sampling was without replacement, the ibd probability for two alleles from one individual is the inbreeding coefficient *F* for that individual, whereas sampling with replacement from one individual has ibd probability (1 + *F*)/2. We defer a more extensive discussion to a subsequent publication.

### Estimation

#### Allele Frequencies

It is allelic identity in state (ibs) that can be observed, rather than identity by descent (ibd) and we now consider how ibs data can provide ibd estimates. We start by considering the frequencies of the various alleles at a locus.

We can distinguish three classes of allele frequency. The sample frequency 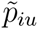 of allele u in a sample of alleles taken from population *i* provides an estimate of the actual allele frequency 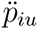 in that population. These actual frequencies, in turn, vary about the population frequency *p*_*u*_, where variation refers to the values of the actual frequencies 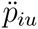 in evolutionary replicates of population *i*.

For *n*_*i*_ randomly sampled alleles, where *n*_*iu*_ are of type *u*, 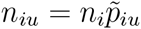 has a binomial distribution 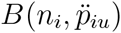. The mean of 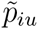 from this statistical sampling process is 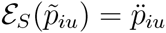 and the variance is 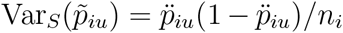. The distribution of the actual frequency 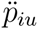 is not known in general, but there is a class of evolutionary models (e.g. Balding and Nichols, 1995) that provide the Beta distribution:

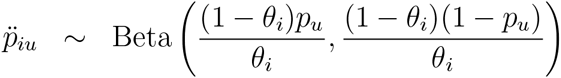

The mean for this genetic sampling process is 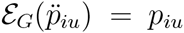 and the variance is 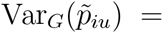 *p*_*u*_(1 − *p*_*u*_)*θ*_*i*_. We keep these first two genetic-sampling moments although we do not invoke the beta distribution.

The total mean and variance follow from

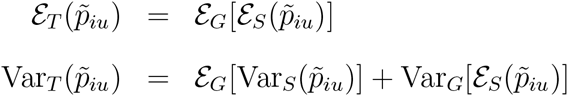

and the complete set of first and second moments were given by Weir and Hill (2002):

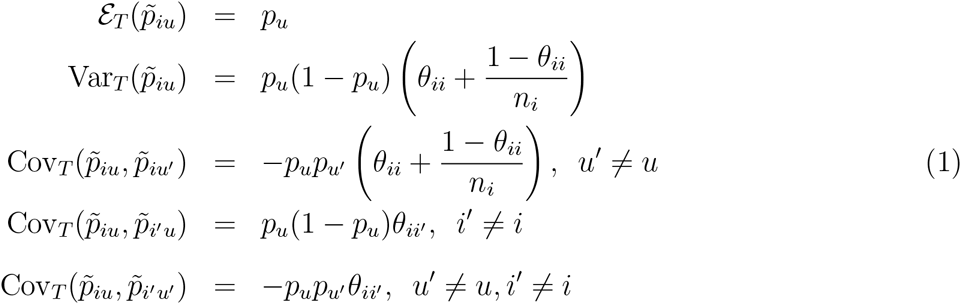

These moments refer to expectations over both repeated samples from the same populations (statistical sampling) and over replications of the populations themselves (genetic sampling). From now on we will drop the *T* subscript but all expectations are total. WC84 set all within-population *θ*_*ii*_ to a common value *θ*, and all between-population *θ*_*ii*′_ to zero. Note the assumption that all populations have the same expected allele frequencies *p*_*u*_, although they have different actual frequencies 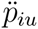.

#### Allelic Matching

We find intuitive appeal in working with proportions of pairs of alleles that are ibs. If the sample of *n*_*i*_ alleles from population *i* has *n*_*iu*_ copies of allele type *u*, then the matching (allele sharing) proportion for pairs of alleles drawn without replacement from population *i* is 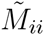 where

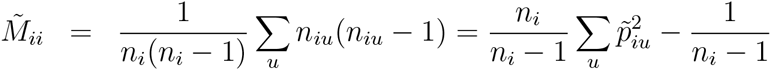

For sampling with replacement the sample-size corrections are not necessary 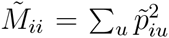. The average over samples from *r* populations is 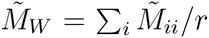. The allele-pair matching proportion between populations *i* and *i*′ is

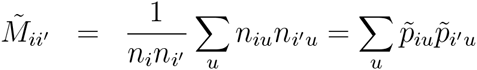

and these have an average over pairs of samples from *r* populations of 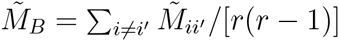.

#### Population Structure Estimates

From Equations 1, the matching proportions have expectations

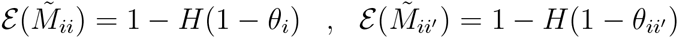

where 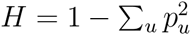. Averaging over populations or pairs of populations:

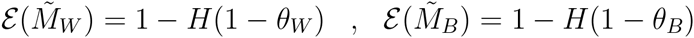

The expectations lead immediately to simple method-of-moment estimates for any number of sampled populations, any number of alleles sampled per population, and any numbers of alleles per locus:

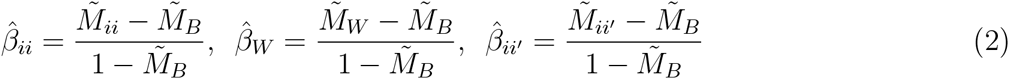

To the extent that the expectation of a ratio is the expectation of ratios, Equations 1,2 show that each 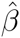 is unbiased for the corresponding *β*:

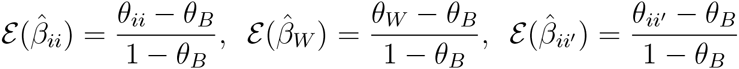

Note that the pairwise estimates 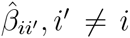 sum to zero by construction. Although it is not possible to find estimates for each *θ* when the sampled populations have correlated sample allele frequencies, it is possible to rank the 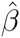’s, and these are likely to have the same ranking as the *θ*’s. We now show how this approach also gives estimates of individual inbreeding coefficients and individual-pair coancestry coefficients.

#### Relatedness Estimates

Suppose now we have a series of individuals *i*; *i* = 1, 2, … *r* and we sample two alleles with replacement from an individual or one allele randomly from each of two individuals. Each individual is regarded as a population with its own actual allele frequencies. Setting the sample sizes to 2 in the matching proportions for one individual and to 1 each for pairs of individuals in the previous section leads to estimates of *θ*_*ii*_ = (1 + *F*_*i*_)/2 or *θ*_*ii*′_ for individual *i* or individuals *i*, *i*′. The expectations of these estimates are [(1 + *F*_*i*_)/2 − *θ*_*B*_]/(1 − *θ*_*B*_) and (*θ*_*ii*′_ − *θ*_*B*_)/(1 − *θ*_*B*_), respectively, where *θ*_*B*_ is the average of all *θ*_*ii*′_, *i* ≠ *i*′. Inbreeding and coancestry are estimated relative to the average coancestry of all pairs of individuals in the study. Yang at al. (2010) also discuss estimates relative to the study population, and say “Estimates of relationships are always relative to an arbitrary base population in which the average relationship is zero. We use the individuals in the sample as the base so that the average relationship between all pairs of individuals is 0 and the average relationship of an individual with him- or herself is 1.” Although our estimates of pairwise relationship sum to zero, we retain the unknown value *θ*_*B*_ in their expectations. We cannot estimate *θ*_*B*_ and we may prefer to report estimates relative to those for the least related pairs as described below in Equation 6.

It is customary (e.g. Yang et al., 2011) use allelic dosage to express relatedness estimates or other analyses (Patterson et al., 2006). Writing the number of copies of allele u carried by individual *i* as *x*_*iu*_, the *U*-allele versions of these standard estimates are

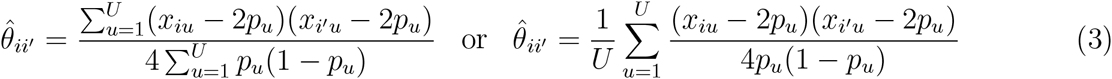

where *i*, *i*′ may be the same or different. If the allele frequencies *p*_*u*_ are known, these estimates are unbiased for *θ*_*ii*′_. For biallelic SNPs, there is no need to sum over alleles, and the *u* subscripts can be dropped.

Our coancestry estimates have the same functional form as those for population structure, but they may best be compared to the standard estimates by expressing allelic matching proportions in terms of allelic dosages. Noting that individual matching proportions are 1 and 0 for homozygotes and heterozygotes, respectively, and that matching proportions for pairs of individuals are 1 when they are the same homozygote, 0.5 when they are the same heterozygote or one is homozygous and the other heterozygous with one allele shared with the first, and 0 when they have no shared alleles:

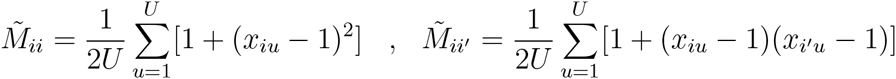

In particular, for SNPs,

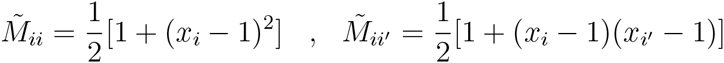

where *x*_*i*_ are the dosages for, say, the reference allele. Our relatedness and inbreeding estimates are

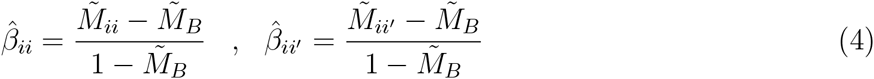

where 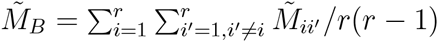.

Storey and Ochoa (accompanying papers) have equivalent estimates. Their expressions are a little different because their reference is for all pairs of alleles in a sample, including those within individuals, whereas ours is for pairs of alleles in different individuals. Astle and Balding (2009, equation 2.3) gave similar estimates although, in effect, they set *θ*_*B*_, the average coancestry of all pairs of individuals in a sample, to zero.

#### Combining Over Loci

Single-locus analyses do not provide meaningful results, and combining estimates over loci *l* has often been considered in the literature. In a parallel discussion of weighting over alleles *u* at a single locus, Ritland (1996) considered weights *w*_*u*_ chosen to minimize variance.

Two extreme weights are *w_l_* = 1 and 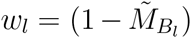. The first may be called “unweighted” and the second “weighted”. In an obvious notation

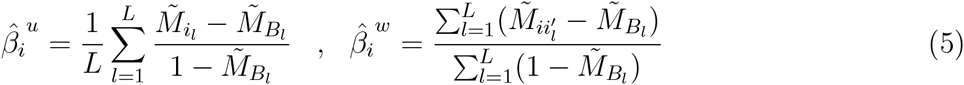

Note the parallel to averaging over alleles in Equations 3. Bhatia et al. (2013) refer to the first estimate as the “average of ratios” and the second as the “ratio of averages.” WC84 advocated the second, with justification given in the Appendix to that paper, as did Bhatia et al.

The unweighted estimate 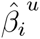 is unbiased for all allele frequencies but is susceptible to the effects of rare variants, when 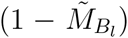 can be very small. Rare variants may have little effect on the weighted average 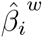, and the variance of the estimate is seen in simulations to be less than for the unweighted average, but it is unbiased only if every locus has the same value of the ibd probabilities. A more extensive discussion was given in the Appendix of WC84 for population structure, and by Ritland (1996) for inbreeding and relatedness.

#### Private Alleles

Current sequence-based studies are revealing large numbers of low-frequency variants, including those found in only one population. These private alleles were identified by Slatkin (1985) and Mathieson and McVean (2012) as being of particular interest. They are very frequent in the 1000 genomes project data (The 1000 Genomes Project Consortium, 2010). If *x*_1_ is the sample count of an allele observed only in population 1 of *r* populations 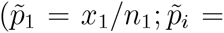 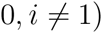 the sample matching proportions are

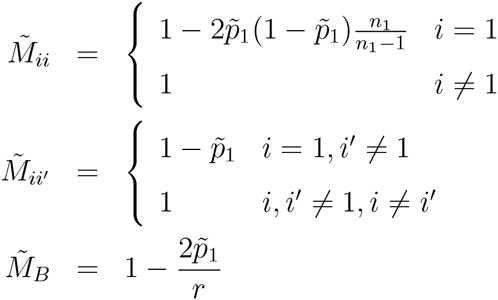

So the *β* estimates are

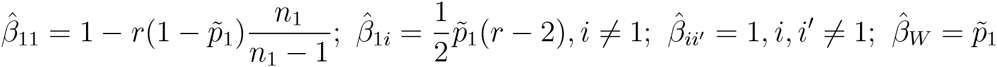

The estimate of *F*_*ST*_ for a private allele is its own-population sample frequency, but the population-specific value for its own population ranges from approximately −*r* + 1 when *x*_1_ = 1 to 1 when *x*_1_ = *n*_1_. This amplifies the comment “populations can display spatial structure in rare variants, even when Wright’s fixation index *F*_*ST*_ is low” of Mathieson and McVean (2012). A population with many private alleles at low to intermediate frequencies will thus likely have a negative 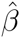, and how negative will depend on how many populations have been sampled. Note that this implies 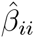 must be allowed to go negative, whereas Bayesian estimators of population specific *F*_*ST*_ are forced to belong to [0, 1], although this assumption can be relaxed (Ritland, 1996).

## RESULTS

### Population Structure

We have conducted a series of simulations to evaluate the performance of our *F*_*ST*_ estimates, and we have looked at 1000 Genomes SNP data to explore the role of rare variants on the estimates. Some of the simulations were conducted with *sim.genot.metapop.t* available in the *hierfstat* package (Goudet, 2005). The migration model we have used allows for a matrix of migration rates between each pair of populations, and the mutation model allows for multiple alleles at a locus. The notation for a two-population model was given above.

#### Model 1. Same Migration Rates, Different Population Sizes

We considered two populations, with sizes *N*_1_ = 100, *N*_2_ = 1, 000 and migration rates *m*_1_ = *m*_2_ = 0.01. The mutation rate was *µ* = 10^−6^. After 400 generations, the *β*’s have values *β*_1_ = 0.156, *β*_2_ = −0.037 and *β*_12_ = 0.059. We simulated 50 individuals from each population under this scenario, with 1,000 loci and up to 20 alleles per locus. From the resulting allelic data we obtained estimates, and 95% confidence intervals by bootstrapping over loci. The results are shown in Table 1. The predicted values are contained in the confidence intervals, and there are negative values for both the parametric and the estimated value of *β*_2_. Note that we cannot estimate *β*_12_ with data from two populations.

**Table 1.** Predicted and estimated *β*’s for two populations

| Model | $N_1$ | $N_2$ | $m_1$ | $m_2$ | $\beta_1$ | $\hat{\beta}_1$ | $\beta_2$ | $\hat{\beta}_2$ | $\beta_{12}$ |
| --- | --- | --- | --- | --- | --- | --- | --- | --- | --- |
| 1 | 100 | 1,000 | 0.01 | 0.01 | 0.156 | 0.163 (0.152, 0.173) | -0.037 | -0.039 (-0.047,-0.032) | 0.059 |
| 2 | 100 | 1,000 | 0.01 | 0.01 | 0.198 | 0.199 (0.192, 0.206) | 0.024 | 0.026 ( 0.023, 0.029) | 0 |
| 3 | 100 | 1,000 | 0.01 | 0.01 | 0.278 | 0.283 (0.268, 0.296) | -0.061 | -0.059 (-0.067,-0.050) | 0.112 |
| 4 | 10,000 | 100 | 0.01 | 0 | -0.319 | -0.302 (-0.409, -0.219) | 0.489 | 0.468 ( 0.372, 0.599) | 0.085 |

Mutation rate $\mu = 10^{-6}$
95% confidence intervals from bootstrapping over loci.

#### Model 2. Continent-Island Model

In this scenario we have an infinite continent supplying a proportion *m* = 0.01 of the alleles independently to populations 1 and 2, still with sizes *N*_1_ = 100, *N*_2_ = 1, 000. There is no migration between the two populations, so *θ*_12_ = 0. The predicted values and estimated values after 400 generations are shown in Table 1.

#### Model 3. Migrant-pool Island Model

In this scenario, each population contributes to a migrant pool, from which migrant alleles are drawn. Among the migrant alleles in the case of two populations, half of the “migrant alleles” will in fact be resident alleles if the gametic pool is composed of the same proportion of alleles from each island, independent of its size. With otherwise the same parameter values, the predicted values and our estimates after 400 generations are shown in Table 1.

#### Model 4. Different Population Sizes, Different Migration Rates

We return to the two-populations model described above, but now with *N*_1_ = 10, 000, *N*_2_ = 100 and different migration rates *m*_1_ = 0.01, *m*_2_ = 0. Predicted values after 1,000 generations are shown in Figure 3, and our estimates in Table 1.

**Figure 3.**
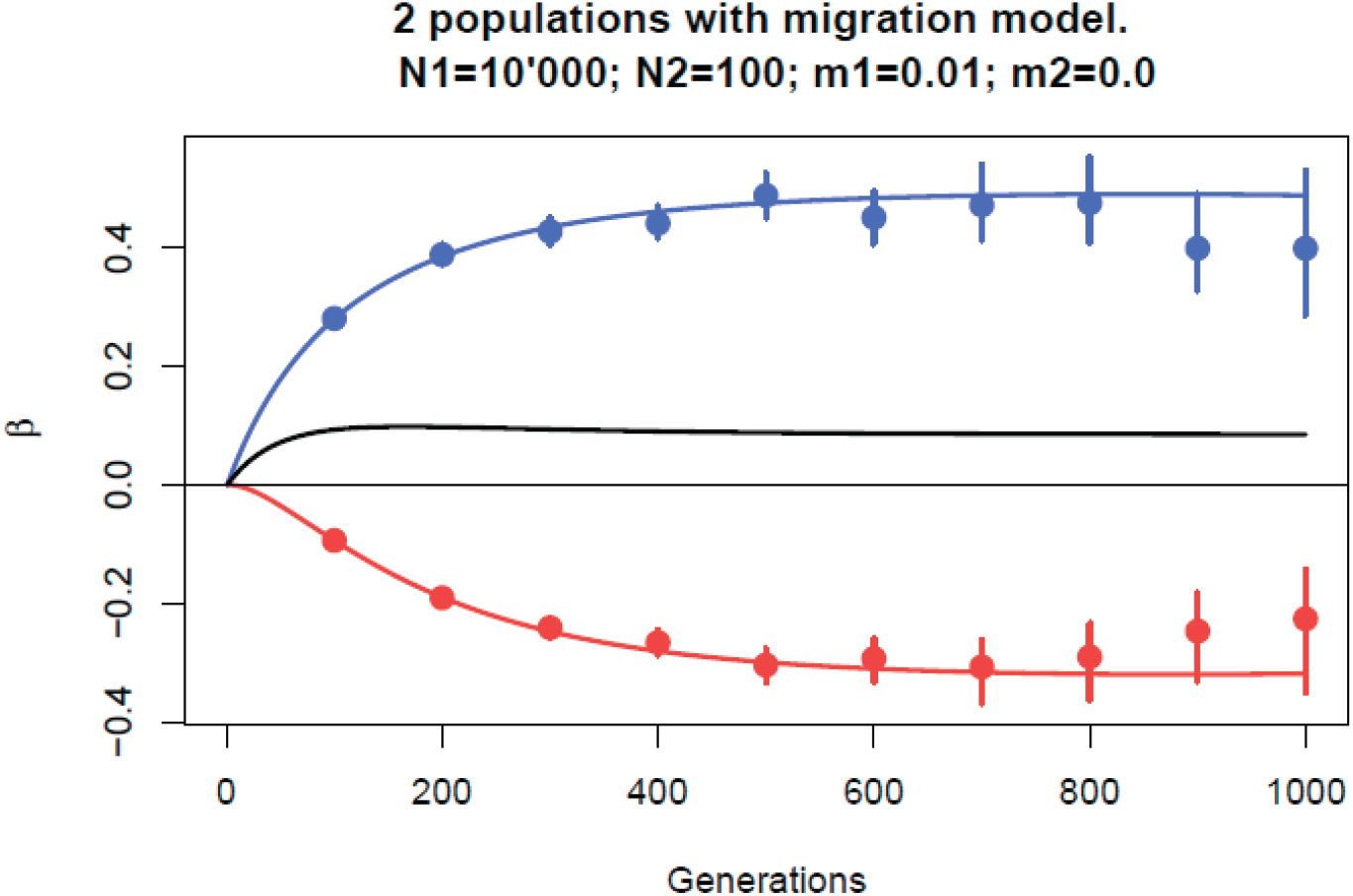
Estimated *β*’s for independent simulations at different times, showing increase in bias and variances as the number of polymorphic loci decreases.

The results in Table 1 show generally good behavior of our *β* estimates. In Figure 3 we show the estimates for 10 different time points (independent replicates) for Model 4. As time increased, the number of polymorphic loci decreased. In generations 600, 800, 1000 the numbers of polymorphic loci had dropped from 1,000 to 712, 349 and 151 respectively and the quality of the estimates decreased: higher bias and higher variance.

#### Rare Alleles

For *r* populations with total sample size *n*_*T*_, and with *x*_1_ copies of an allele private to population 1, the total count for this alleles is *x*_*T*_ = *x*_1_ and 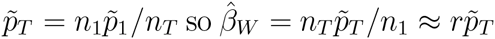 assuming similar sample sizes for each sample. In Figure 4 we display 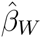 as a function of allele frequencies for SNPs located on chromosome 2 in the 1000 Genomes project. Individuals were grouped by regions (Africa, Europe, South Asia, East Asia and the Americas). The drawn line corresponds to *β*_*W*_ = 5*p*_*T*_. The initial linear segment corresponds to alleles that are present in one continent only. *β*_*W*_’s start departing from this line for allele counts larger than 80, or equivalently, for worldwide frequencies larger than ≈ 0.01, given the sampled chromosome number of 2, 426.

**Figure 4.**
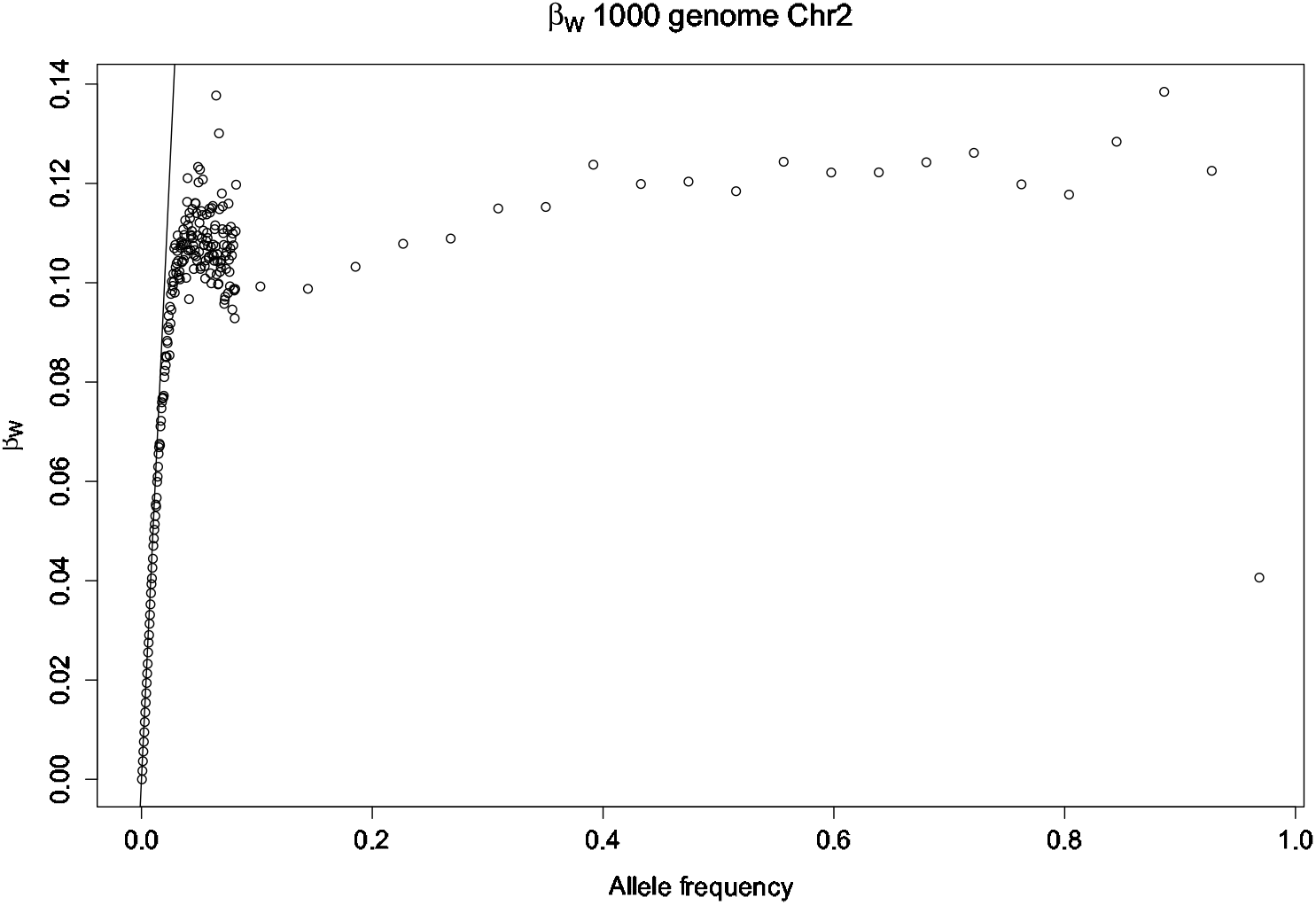
*β*_*W*_ as a function of allele frequencies (*n*_*u*_ /*n_T_*) for SNPs located on chromosome 2. Data from the 1000 genomes project, individuals were grouped by regions (Africa, Europe, South Asia, East Asia and Americas). The drawn line corresponds to 5*n*_*u*_ /*n_T_*. The initial linear segment corresponds to alleles that are present in one continent only. *β*_*W*_ s start departing from this line for allele counts larger than 80, or equivalently, for worldwide frequencies larger than ≈ 0.01, given the sampled chromosome number of 2426.

When a new allele appears, it will be present in one population only. We expect most if not all rare alleles to be private alleles, and thus the expected values for *F*_*ST*_ (*β*_*W*_) for these rare alleles are their (sub-population) frequencies. When *β*_*W*_ starts departing from the allele frequency, it implies that some scattering has been happening. In species with a lot of migration, this will happen at low frequencies, whereas the species that are more sedentary should show a one to one relation between sub-population allele frequencies and *β*_*W*_ for a larger range of their site frequency spectrum.

In Buckleton et al. (2016) we gave population-specific *F*_*ST*_ estimates for a set of 446 populations, using published data for 24 microsatellite loci collected for forensic purposes. We showed in that paper how the choice of a reference set of populations can affect results. For a set of African populations, the average within-population matching proportion was 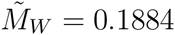 and the average between-population-pair averages were 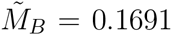 within the African region and 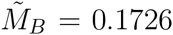 for all pairs of populations. There is a larger *F*_*ST*_ for the set of African populations 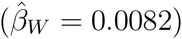 with Africa as a reference set than there is 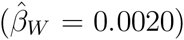 with the world as a reference set. The opposite was found for a collection of Inuit populations: the average within-population matching proportion was 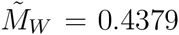 whereas the average between-population-pair matching proportions were 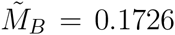 for pairs within the Inuit group and 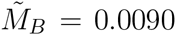 for all pairs in the study: so *F*_*ST*_ is less with Inuit as a reference set 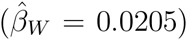 than with the world as a reference set 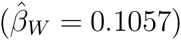.

### Inbreeding and Relatedness

To check on the validity of our estimators of individual inbreeding and coancestry coefficients, we simulated data for a range of 11 coancestries: (*i*/32 : *i* = 0, 1, 2, …, 10). Using the *ms* software (Hudson, 2002), we generated data from an island model with two populations exchanging *N m* = 1 migrant per generation. We simulated 5,000 independent loci, read either as haplotypes (5,000) or as SNPs (approximately 80,000 polymorphic sites for the founders). We then chose 20 individuals from one of these populations and let them mate at random, without selfing. We did not assign or consider sex for these 20 founders. The number of offspring per mating was Poisson with mean of five. These offspring were than allowed to mate at random, without selfing, to produce families of size Poisson with mean three. By keeping records of all matings we could generate the pedigree-based inbreeding and coancestry values for all 135 individuals: founder, their offspring and their grand-offspring. The pedigree-based coancestries for all 9,045 pairs of individuals are shown in Figure 5, although we note (Hill and Weir, 2011) that the actual values have variation about expected or pedigree values. We used the same pedigree to simulate another data set, where this time 10 founders were coming from the first population and the other 10 from the second population, thus creating admixture among the children and grand-children.

**Figure 5.**
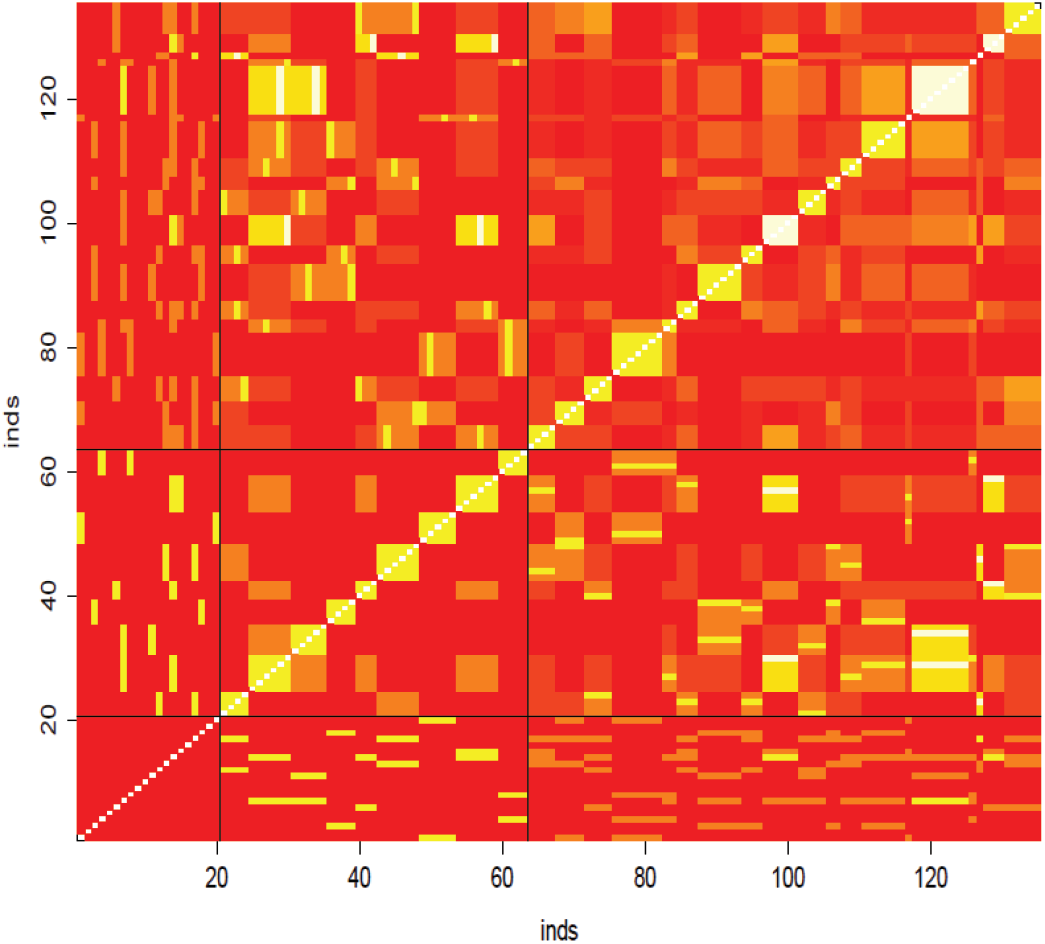
Pedigree-based inbreeding and coancestry coefficients for simulated data.

The left hand plot of Figure 6 reflects the summing to zero by construction of the 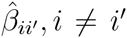 coancestries, whereas the pedigree coancestries are necessarily non-negative. The right hand plot shows a “correction” of the estimates: we took the set of smallest 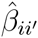 values in the left hand plot to represent the unrelated (relative to the assumed-unrelated) founders. If we write 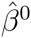 as the average value in this distribution then our corrected values 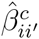 are

**Figure 6.**
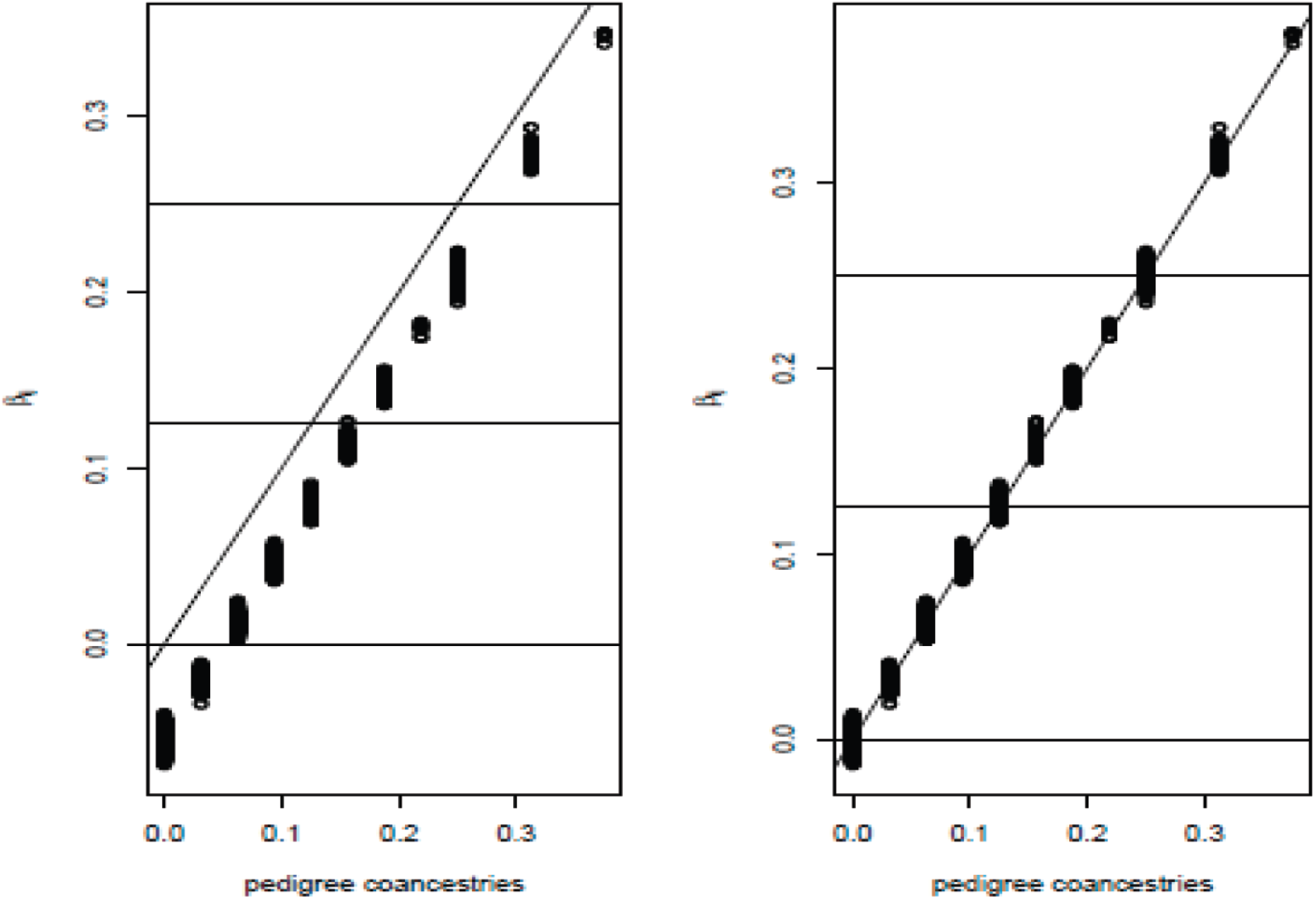
Comparison of estimated and pedigree coancestries. Uncorrected estimates on left, corrected estimates on right. Correction procedure described in text.

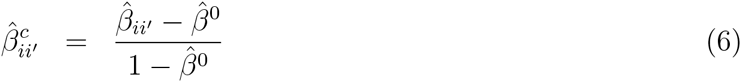

The corrected estimates are clearly close to the pedigree values. However, we are not sure if it is necessary, in general, to undertake this correction process. Whether or not it is applied, the 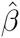 values are still relative to those among all pairs of individuals in a study sample. In general, we will not have any individuals identified for which it is justified to assume zero relatedness or zero inbreeding, and we note the comment by Thompson (2013) “in most populations IBD within individuals is at least as great as IBD between.”

The distributions of estimates in Figure 7 are tightly clustered around 11 values, corresponding to the 11 distinct pedigree values *i*/16, *i* = 0, 1, 2 … 11. A contrasting result is shown in Figure 8, for the CGTA estimates, calculated as weighted averages over loci (in the sense of equation 3 by taking the ratio of two sums over loci).

**Figure 7.**
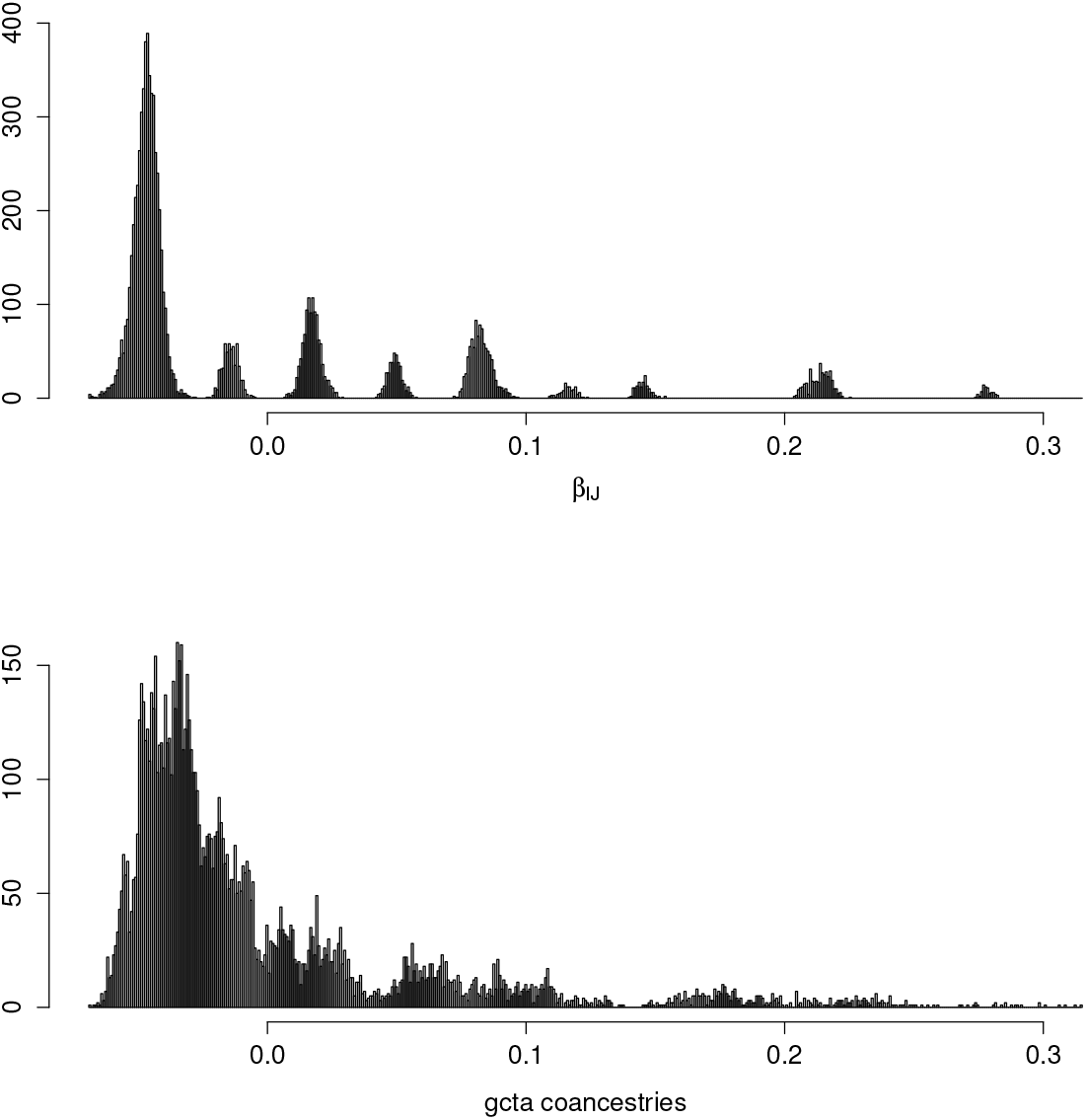
Comparison of *β* and CGTA coancestry estimates, when founders are drawn from a single population.

**Figure 8.**
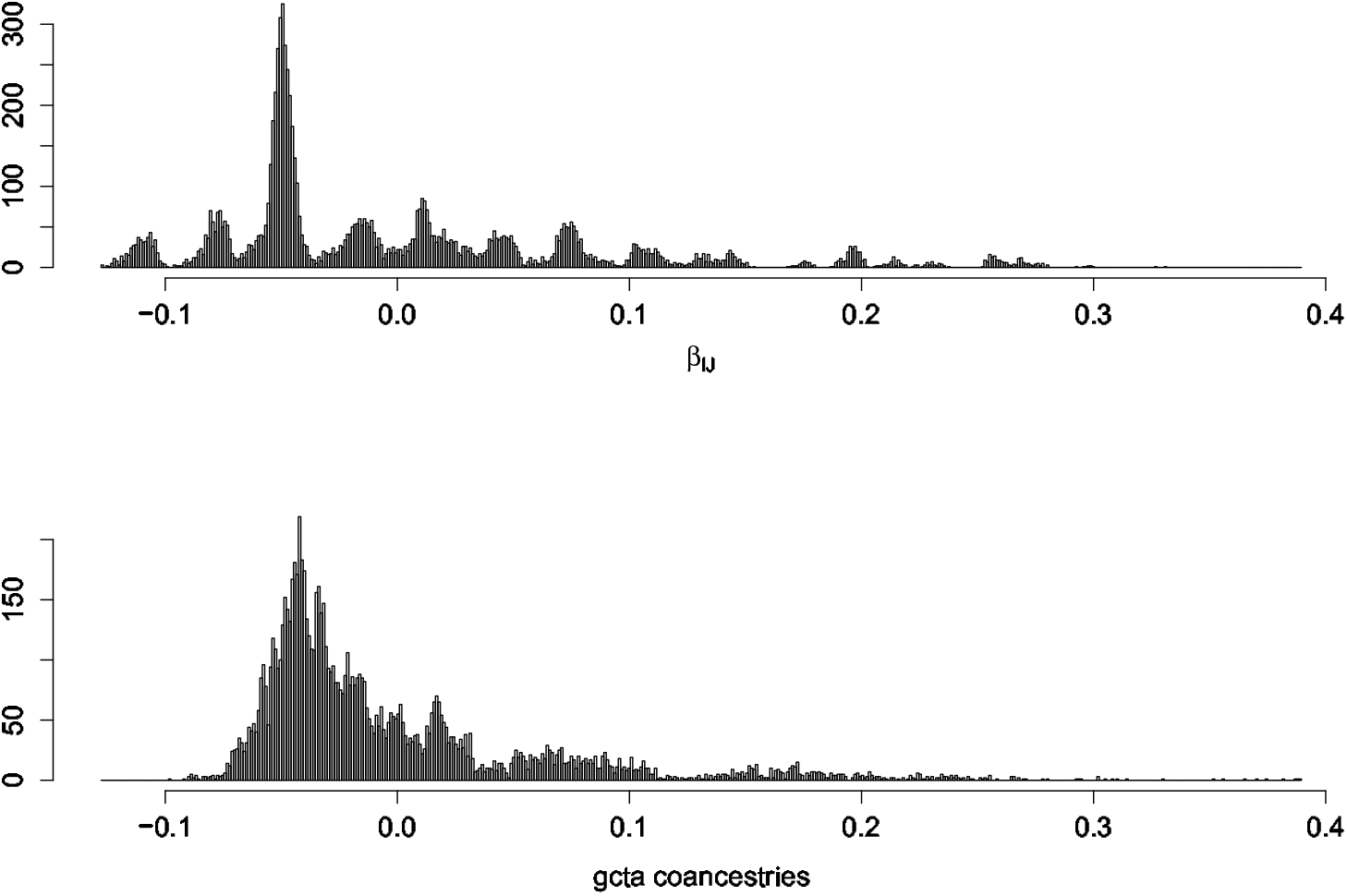
Comparison of *β* and CGTA coancestry estimates, when founders are drawn from two populations.

There is a current tendency in genome wide association studies (GWAS) to restrict the SNPs used in relatedness estimation to having a minor allele frequency (MAF) above some threshold. For example, the KING manual (http://people.virginia.edu/∼wc9c/KING/manual.html) lists a parameter **minMAF** to specify the minimum minor allele frequency to select SNPs for relationship inference in homogeneous populations. The thought is that lesser frequencies give rise to biased values, but that is not likely the case if “ratio of averages” estimates are used. To illustrate the effect of MAF filtering, we applied four different thresholds for our simulated data and we show the means and standard deviations for estimates for each of nine pedigree values in Table 2. The estimates are the corrected values – i.e. relative to an assigned value of zero for the least-related class. There is clear evidence for the merits of retaining all SNPs, both in terms of bias and variance.

**Table 2.** Effects of filtering to L SNPs on coancestry estimate means (and standard deviations ×100).

| | $L = 79,069$ | $L = 72,012$ | $L = 56,979$ | $L = 44,061$ |
| --- | --- | --- | --- | --- |
| Pedigree value | All SNPs | MAF $\geq 0.01$ | MAF $\geq 0.05$ | MAF $\geq 0.10$ |
| 0 | 0.000 (0.50) | 0.000 (1.00) | 0.000 (1.99) | 0.000 (2.43) |
| 0.03125 | 0.031 (0.30) | 0.026 (0.30) | 0.010 (0.89) | 0.003 (1.45) |
| 0.06750 | 0.061 (0.34) | 0.056 (0.35) | 0.041 (1.13) | 0.036 (1.79) |
| 0.09375 | 0.092 (0.27) | 0.087 (0.27) | 0.069 (0.72) | 0.061 (1.13) |
| 0.12500 | 0.124 (0.41) | 0.120 (0.46) | 0.112 (1.90) | 0.109 (2.69) |
| 0.15625 | 0.156 (0.29) | 0.151 (0.29) | 0.133 (0.65) | 0.122 (1.15) |
| 0.18750 | 0.184 (0.26) | 0.179 (0.27) | 0.157 (1.07) | 0.144 (1.64) |
| 0.25000 | 0.249 (0.42) | 0.245 (0.45) | 0.241 (1.87) | 0.239 (2.62) |
| 0.31250 | 0.311 (0.20) | 0.307 (0.20) | 0.285 (0.77) | 0.271 (1.23) |

We continued a comparison of 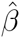 values by applying the estimates described by Wang (2014) and computed using the *related* R package (Pew et al., 2015), listed in Table 3. Additionally, *related* offers maximum likelihood estimators, derived by Milligan (2003) and Wang and Santure (2007). They are not computed here, because they require substantial computing time, which rules them out for genomic data.

**Table 3.** Other estimates of relatedness.

| Method | Description |
| --- | --- |
| `ped` | The pedigree based relatedness. It can be considered as the true relatedness value, although it will depend on the depth of the pedigree. |
| `bij` | $\beta_{ij}$, developed here. These values are relative to the mean of the population, and hence, the mean of these relatedness must be 0. |
| `wang` | The estimator developed by Wang (2002). |
| `lynchli` | The estimator derived by Lynch (1988) and improved by Li et al. (1993), equation (7) in Wang (2014). |
| `lynchrd` | The estimator derived by Lynch and Ritland (1999) [eq (5,6) in Wang, 2014]. |
| `GCTA` | The estimator used in GCTA (Yang et al., 2011). For multi-allelic markers, alleles are weighted according to their variance. |
| `ritland` | The estimator derived by Ritland (1996) [eq (4) in Wang (2014)]. For SNPs, it is the same as `GCTA`, but for multi-allelic markers, each allele is given the same weight. |
| `quellergt` | The estimator derived by Queller and Goodnight (1988) [eq (2,3) in Wang (2014)]. |

In Figure 9 we display box plots of coancestry estimates for eight alternative estimates, displayed according to 9 pedigree values. The *β* estimates are not corrected, yet have good bias and variance properties compared to other estimates.

**Figure 9.**
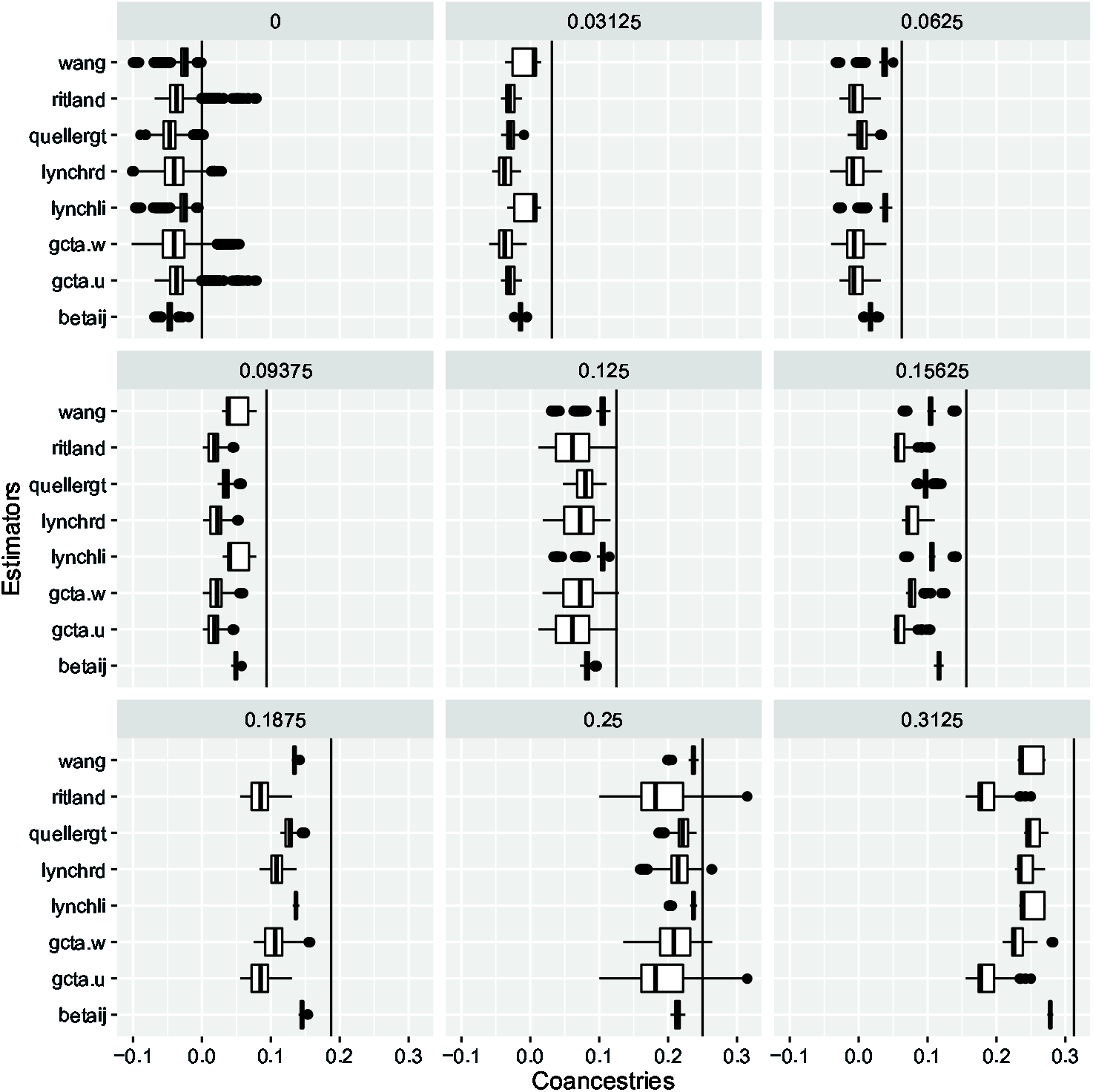
Boxplots of coancestry estimates for eight alternative estimates, displayed according to nine pedigree values. The *β* estimates are not corrected, yet have good bias and variance properties.

In Figure 10 we compare our *β* estimates with those from GCTA for admixed individuals with two ancestral populations. We used the same pedigree as in the section above, but took as founders 10 individuals from each of the two populations. Coancestries were calculated for all pairs of individuals in the pedigree. Figure 10 illustrates the accuracy of our *β* estimate compared with GCTA using the coancestries among founders. The 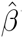’s for pairs of founders from the same population are tightly distributed around 0.015, while 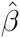’s for pairs of individuals one from each population are tightly distributed around −0.11. The distribution for the same two categories for the GCTA estimators is wider, in particular for pairs of individuals originating from the same population.

**Fig 10.**
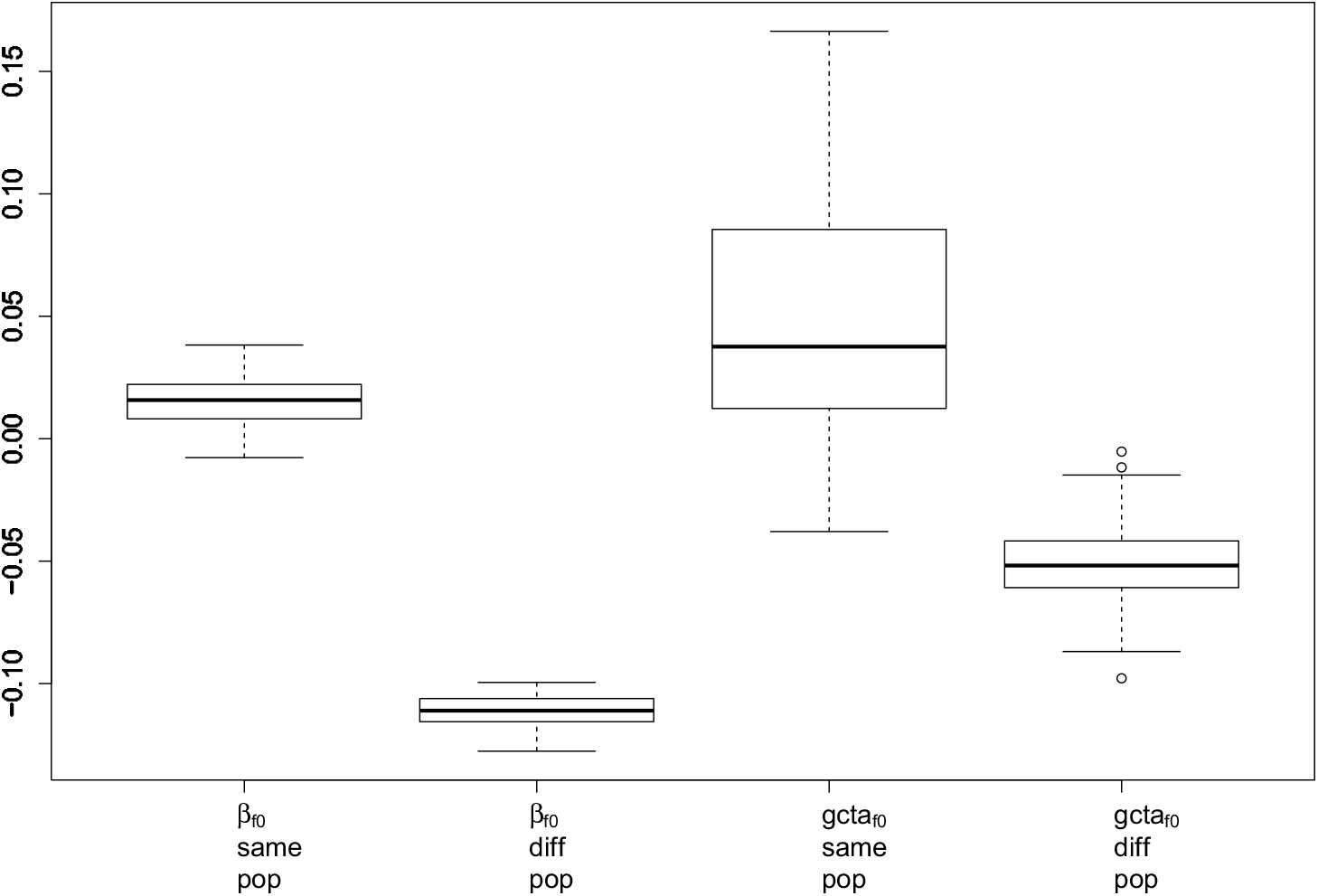
Boxplots of coancestry estimates *β* and GCTA when the founders come from two populations. Coancestries were estimated for all the individuals in the pedigree, but only those between founders are shown. For each panel, the left boxplot is for pairs of founders from the same population, while the right boxplot is for pairs when the two members come from different populations.

## DISCUSSION

### A Unified Approach

Although there has been general recognition that family and evolutionary relatedness are just two ends of a continuum, we are not aware of previous estimates of population structure quantities such as *F*_*ST*_ or individual-pair coancestries that rest on this common framework. We have presented estimates that apply equally well to populations and individuals. While their statistical properties remain to be fully explored, it is reassuring to see how well they performed in the few simulations presented here.

Although individual-specific inbreeding coefficients and individual-pair-specific coancestry coefficients are used routinely in association studies, we have not seen widespread adoption of population-specific *F*_*ST*_ values in evolutionary studies. We have shown here, theoretically and empirically, that these values can differ substantially among populations. This may simply reflect population size and migration rate differences, but different values may also provide signatures of natural selection.

There is also general understanding that identity by descent is a relative concept, rather than an absolute concept. This understanding has not led to an apparent recognition that the usual estimates of inbreeding and kinship are not unbiased for expected or pedigree values. Replacing population allele frequencies by sample values leads to bias in the usual estimates, *regardless of sample size*. As the allele frequencies enter GCTA estimates, for example, as squares the expected values of the estimates depend on the variances these frequencies. These, in turn, depend on the parameters being estimated.

We also stress that all allelic variants, whatever their frequencies, need to be included in the estimation of population structure and inbreeding or relatedness. The estimates certainly depend on the allele frequencies, and restricting the range of frequencies used may reveal features of interest, but the underlying ibd parameters do not depend on the frequencies. Exclusion of some alleles based on their frequencies will lead to biased estimates of the parameters.

### Previous Estimates

#### Weir and Cockerham Estimates of *F*_*ST*_

The *F*_*ST*_ estimate of WC84 has been widely adopted and it performs well for the model stated in that paper: data from a series of independent populations with equivalent histories. In the present notation, WC84 assumed *θ*_*ii*_ = *θ*, *θ*_*ii*′_ = 0 for all populations i and all *i*′ ≠ *i*. The estimate was designed to be unbiased for any number of populations, any sample sizes and any number of alleles per locus. The analysis was a weighted one over population: the average allele frequencies 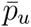 for a study had sample size weights, 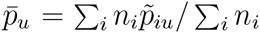. Although our *β* estimates do not make explicit mention of allele frequencies, there is implicit use of sample frequencies that are unweighted averages over individuals or populations.

Weighting over populations has been discussed by Tukey (1957) and Robertson (1962). Those authors were concerned with bias and variance and they used the language of variance components, within and between populations. For allele u these components were given as (1 − *θ*)*p*_*u*_ (1 − *p*_*u*_) and *θp*_*u*_ (1 − *p*_*u*_), respectively, by WC84. Tukey said “In practice, we select two quadratic functions by some scheme involving intuition, find how their average values are expressed linearly in terms of the variance components, and then form two linear combinations of the original quadratics whose average values are the variance components. These linear combinations are then our estimates. Much flexibility is possible.” The estimates of WC84, Weir and Hill (2002) and Bhatia et al. (2013) all have this structure although ratios of linear combinations are taken to remove the allele frequency parameters. Tukey went on to say that the weights *w*_*i*_ = *n*_*i*_ (in the present notation) “gives the customary analyses, which treat observations as important and columns [i.e. populations] as unimportant.” Further, “the choice *w*_*i*_ = 1 … treat the columns as important. This [unweighted] approach is appropriate when the column variance component is large compared with the within variance component.” Robertson (1962) also pointed to sample-size weights for small between-population variance components and equal weights for large values. Bhatia et al. (2013) were concerned with unequal *F*_*ST*_ values so their use of equal weights is consistent with Turkey’s statements. Their work provides simple averages of the different *F*_*ST*_’s as opposed to averages weighted by sample sizes. For unequal *F*_*ST*_’s and unequal sample sizes, Weir and Hill (2002) said “the usual moment estimate [with sample-size weights] is of a complex function [of the *F*_*ST*_’s].” In our current model of unequal *θ*_*i*_’s and non-zero *θ*_*ii*′_’s we agree that unweighted analyses (population weights of 1) are appropriate, and that is what we have used in this paper. We note that Tukey’s “flexibility” in the choice of moment estimators, phrased in terms of weights, does not arise with maximum likelihood approaches. If sample allele frequencies are taken to be approximately normally distributed then REML methods give appropriate and unique estimates.

What are the consequences of using the WC84 estimates when the current model is more appropriate? We can show that the expected value of the Weir and Cockerham estimate 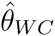 is

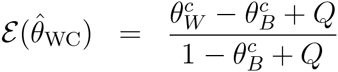

This expression uses three functions of sample sizes: 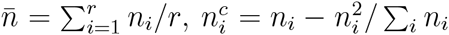 and *n*_*c*_ = 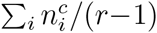. The two weighted averages are 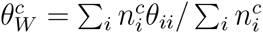 and 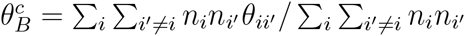. The quantity *Q* is 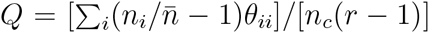. For equal sample sizes, *n*_*i*_ = *n*, or for equal values of *F*_*ST*_, 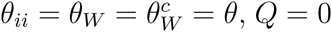. Under these circumstances 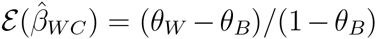 and we find the WC84 estimator performs well unless the *θ*_*ii*_ ‘s and /or the *n*_*i*_’s are quite different. We stress though that it is (*θ*_*W*_ – *θ*_*B*_)/(1 – *θ*_*B*_) being estimated.

#### Nei Estimates of *F*_*ST*_

Although we have phrased estimates in terms of matching proportions, we note that they are the complements of “heterozygosities” 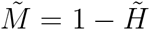. Our approach uses 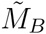, the average population-pair allele matching, whereas most previous treatments, from Nei (1973) onwards, use total heterozygosities 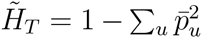 where 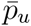 is the average sample allele frequency over populations: 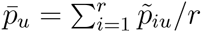. From Equations 1, the variance of 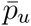 is

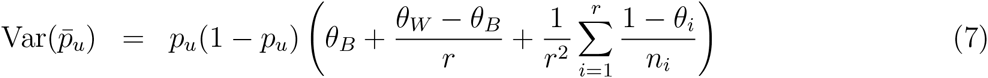

For large sample sizes 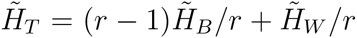 and Nei’s *G*_*ST*_ quantity and its expectation, in our notation, are

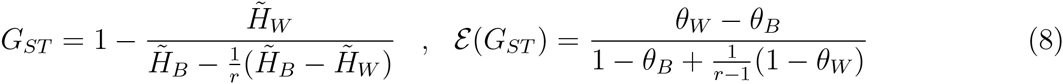

which reduce to 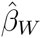 and 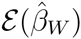 as *r* becomes large. Otherwise, the expectation of *G*_*ST*_ depends on the number *r* of populations. This expectation is bounded above by one, contrary to the claim of Bhatia et al. (2013). Bounds on *F*_*ST*_, when that is defined as 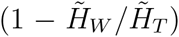, were given by Jakobsson et al. (2013).

Nei and Chesser (1983) and Nei (1987) modified Nei’s earlier approach to remove the effects of the number of populations. Jost (2008) pointed out that *G*_*ST*_ does not provide a good measure of differentiation among populations, where differentiation reflects the collection of allele frequencies 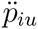, or their sample values 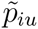. We regard *θ*’s as indicators of evolutionary history, rather than of allele frequencies, and we interpret them as probabilities of pairs of alleles being identical by descent. Jost introduced *D* = (*H*_*B*_ − *H*_*W*_)/(1 − *H*_*W*_) or *D* = (*θ*_*W*_ − *θ*_*B*_)/*θ*_*W*_ as a measure of differentiation among populations. For the two-population drift scenario without mutation *D*, unlike *β*_*W*_, does not have a simple dependence on time and so does not serve as a measure of evolutionary distance.

#### CGTA Estimates of Relatedness

The expressions in Equation 3 provide unbiased estimates of *θ*_*ii*_ = (1 + *F*_*i*_)/2 and *θ*_*ii*′_, *i* ≠ *i*′ when the allele frequencies are known. When study sample allele frequencies are used the expectations of these expressions, for one locus and large samples, are

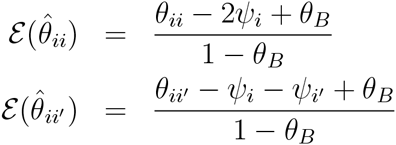

where 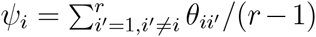. The extent of bias depends on how different the average coancestry of a target individual with all other study individuals is from the average coancestry of all pairs of study individuals. We stress that these estimates are not unbiased for *θ*_*ii*′_.

### Association Mapping

One of our motivations for seeking a unified characterization of population structure and relatedness is that both phenomena affect samples used in association mapping. Many analyses, such as those in GCTA, use mixed linear models with an estimated Genetic Relatedness Matrix **A** being used in the formulation of the variance-covariance matrix for trait values of the study individuals. For a trait with additive genetic variance 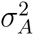 and no other genetic variance components, the variance matrix for individuals includes the term 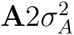 and **A** has diagonal elements (1 + *F*_*i*_)/2 and off-diagonal elements *θ*_*ii*′_. We suggest that these be estimated by 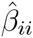 and 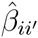, to accommodate any (hidden) relatedness and inbreeding among study subjects. We are less sure about the common practice of also using principal components of **A** as fixed effects to accommodate population structure, especially when **A** uses 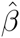’s, and we see the need for further investigation.

### Population History

We also see the need for further exploration of the role of population-specific *F*_*ST*_ estimates in evolutionary genetic studies, given the generally unrecognized prevalence of negative expected values for populations with correlated allele frequencies shown in Figure 1 and the relationship of estimates with the site-frequency spectrum suggested in Figure 4.

## Conclusion

We have presented moment estimators for the probabilities that pairs of alleles, taken from individuals or from populations, are identical by descent relative to the ibd probabilities for alleles from all pairs of individuals or populations in a study. By identifying the reference set of alleles as those in the current study we allow for negative values for population structure or relatedness parameters and their estimates. Alleles may have smaller ibd probabilities within some populations than between all pairs of populations in a study, for example. Some pairs of individuals in a study may be less related than the average for all pairs. Our estimates are phrased in terms of the proportions of pairs of alleles, within and between populations or individuals, that are of the same type (ibs).

For sets of populations, we advocate the use of population-specific *F*_*ST*_ values as these more accurately reflect population history. For sets of individuals, our estimates seem to behave at least as well as those given previously. We note that our estimates have the same logical basis, and algebraic expressions, for populations and for individuals. The chief novelty of our approach is in allowing for allele frequencies to be correlated among populations when characterizing population structure, and correlated among all individuals when characterizing individual-pair relatedness.

## Acknowledgments

This work was supported in part by grants GM 075091 and GM 099568 from the US National Institutes of Health and by grant IZK0Z3 157867 from the Swiss National Science Foundation.

## Literature Cited

Astle W, Balding DJ. 2009. Population structure and cryptic relatedness in genetic association studies. Statistical Science 24:451–471.

Balding DJ, Nichols RA. 1995. A method for quantifying differentiation between populations at multi-allelic loci and its implications for investigating identity and paternity. Genetica 96:3–12.

Beaumont MA, Balding DJ. 2004. Identifying adaptive genetic divergence among populations from genome scans. Molecular Ecology 13:969–980.

Bhatia G, Patterson N, Sankararaman S, Price AL. 2013. Estimating and interpreting *F_ST_*: The impact of rare variants. Genome Research 23:1514–1521.

Browning SR, Weir BS. 2010. Population structure with localized haplotype clusters. Genetics 185:1337–l1344.

Buckleton JS, Curran JM, Goudet J, Taylor D, Thiery A, Weir BS. 2016. Population-specific *F*_*ST*_ values: A worldwide survey. Forensic Science International: Genetics 23:91–100.

Cockerham CC, Weir BS. 1983. Variance of actual inbreeding. Theoretical Population Biology 23:85–109.

Foll M, Gaggiotti O. 2006. Identifying the environmental factors that determine the genetic structure of populations. Genetics 174:875–891.

Gaggiotti OE, Foll M. 2010. Quantifying population structure using the *F*-model. Molecular Ecology Resources 10:821–830.

Goudet J. 2005. hierfstat, a package for R to compute and test hierarchical *F*-statistics. Molecular Ecology Notes 5:184–186.

Hill WG, Weir BS. 2011. Variation in actual relationship as a consequence of Mendelian sampling and linkage. Genetics Research 93:47–74.

Hill WG, Weir BS. 2012. Variation in actual relationship among descendants of inbred individuals. Genetics Research 94:267–274.

Hudson RR. 2002. Generating samples under a Wright-Fisher neutral model. Bioinformatics 18:337–8

Jakobsson M, Edge MD, Rosenberg NA. 2013. The relationship between *F*_*ST*_ and the frequency of the most frequent allele. Genetics 193:515–528.

Jost L. 2008. G(ST) and its relatives do not measure differentiation. Molecular Ecology 17:4015–4026.

Kang HM, Sul JH, Service SK, Zaitlen NA, Kong SY, Freimer NB, Sabatti C, Eskin E. 2010. Variance component model to account for sample structure in genome-wide association studies. Nature Genetics 42:348–354.

Li CC, Weeks DE, Chakravarti, A. 1993. Similarity of DNA fingerprints due to chance and relatedness. Human Heredity 43: 45–52.

Lynch M. 1988. Estimation of relatedness by DNA fingerprinting. Molecular Biology and Evolution 5: 584–599.

Lynch M, Ritland K. 1999. Estimation of pairwise relatedness with molecular markers. Genetics 152: 1753–1766.

Manichaikul A, Mychaleckyj JC, Rich SS, Daly K, Sal M, Chen W-M. 2010. Robust relationship inference in genome-wide association studies. Bioinformatics 26:2867–2873.

Mathieson I, McVean G. 2012. Differential confounding of rare and common variants in spatially structured populations. Nature Genetics 44:243–248.

Maruyama T. 1970. Effective number of alleles in a subdivided population. Theoretical Population Biology 1:27–306.

McTavish EJ, Hillis DM. 2015. How do SNP ascertainment schemes and population demographics affect inferences about population history? BMC Genomics 16:266–278.

Milligan BG. 2003. Maximum-likelihood estimation of relatedness. Genetics 163:1153–1167.

Nei M. 1973. Analysis of gene diversity in subdivided populations. Proceedings of the National Academy of Sciences, USA 70:3321–3323.

Nei M. 1987. Molecular Evolutionary Genetics. Columbia University Press, New York.

Nei M, Chesser RK. 1983. Estimation of fixation indices and gene diversities. Annals of Human Genetics 47:253–259.

Patterson N, Price AL, Reich D. 2006. Population structure and eigenanalysis. PLoS Genetics 2:2074–2093.

Pew J, Muir PH, Wang J, Frasier TR. 2015. related: an R package for analysing pairwise relatedness from codominant molecular markers. Mol Ecol Resour 15: 557–561

Purcell S, Neale B, Todd-Brown K, Thomas L, Ferreira MAR, Bender D, Maller J, Sklar P, de Bakker PIW, Daly MJ, Sham PC. 2007. PLINK: A toll set for while-genome association and population-based linkage analysis. American Journal of Human Genetics 81:559–575.

Queller DC, Goodnight KF. 1989. Estimating relatedness using molecular markers. Evolution 43: 258–275.

Reynolds J, Weir BS, Cockerham CC. 1983. Estimation of the coancestry coefficient: basis for a short-term genetic distance. Genetics 105:767–779.

Ritland K. 1996. Estimators for pairwise relatedness and individual inbreeding coefficients. Genetical Research 67:175–185.

Robertson A. 1962. Weighting in the estimation of variance components in the unbalanced single classification. Biometrics 18:3–17.

Shriver MD, Kennedy GC, Parra EJ, Lawson HA, Sonpa V, Huang J, Akey JM, Jones KW. 2004. The genomic distribution of population substructure in four populations using 8,525 autosomal SNPs. Human Genomics 41:274–286.

Slatkin M. 1985. Rare alleles as indicators or gene flow. Evolution 39:53–65.

The 1000 Genomes Project Consortium. 2010. A map of human genome variation from population-scale sequencing. Nature 467:1061–1073.

Thompson EA. 1975. Estimation of pairwise relationships. Annals of Human Genetics 39:173–188.

Thompson EA. 2013. Identity by descent: variation in meiosis, across genomes, and in populations. Genetics 194:301–326.

Tukey JW. 1957. Variances of variance components: II. The unbalanced single classification. Annals of Mathematical Statistics 28: 43–56.

Wang J. 2002. An estimator for pairwise relatedness using molecular markers. Genetics 160: 1203–1215.

Wang J. 2014. marker-based estimates of relatedness and inbreeding coefficients: an assessment of current methods. Journal of Evolutionary Biology 27:518–530.

Wang J, Santure AW. 2009. Parentage and sibship inference from multilocus genotype data under polygamy. Genetics 181:1579–1594

Weir BS. 1996. Genetic Data Analysis II. Sinauer, Sunderland, MA.

Weir BS, Cardon LR, Anderson AD, Nielsen DM, Hill WG. 2005. Measures of human population structure show heterogeneity among genomic regions. Genome Research 15:1468–1476.

Weir BS, Cockerham CC. 1984. Estimating F-statistics for the analysis of population structure. Evolution 38:1358–1370.

Weir BS, Hill WG. 2002. Estimating F-statistics. Annual Review of Genetics 36:721–750.

Yang J, Benyamin B, McEvoy BP, Gordon S, Henders AK, Nyholt DR, Madden PA, Heath AC, Martin NG, Montgomery GW, Goddard ME, Visscher PM. 2010. Common SNPs explain a large proportion of the heritability for human height. Nature Genetics 42:565–569.

Yang J, Lee SH, Goddard ME, Visscher PM. 2011. GCTA: A tool for genome-wide complex trait analysis. American Journal of Human Genetics 88:76–82.

Yu J, Pressoir G, Briggs WR, et al. 2006. A unified mixed-model method for association mapping that accounts for multiple levels of relatedness. Nature Genetics 38:203–208.

